# Transparent exploration of machine learning for biomarker discovery from proteomics and omics data

**DOI:** 10.1101/2021.03.05.434053

**Authors:** Furkan M. Torun, Sebastian Virreira Winter, Sophia Doll, Felix M. Riese, Artem Vorobyev, Johannes B. Mueller-Reif, Philipp E. Geyer, Maximilian T. Strauss

**Affiliations:** OmicEra Diagnostics GmbH, Planegg, Germany

**Keywords:** Artificial intelligence, mass spectrometry, diagnostics, omic, proteome, metabolome, transcriptome

## Abstract

Biomarkers are of central importance for assessing the health state and to guide medical interventions and their efficacy, but they are lacking for most diseases. Mass spectrometry (MS)-based proteomics is a powerful technology for biomarker discovery, but requires sophisticated bioinformatics to identify robust patterns. Machine learning (ML) has become indispensable for this purpose, however, it is sometimes applied in an opaque manner, generally requires expert knowledge and complex and expensive software. To enable easy access to ML for biomarker discovery without any programming or bioinformatic skills, we developed ‘OmicLearn’ (https://OmicLearn.com), an open-source web-based ML tool using the latest advances in the Python ML ecosystem. We host a web server for the exploration of the researcher’s results that can readily be cloned for internal use. Output tables from proteomics experiments are easily uploaded to the central or a local webserver. OmicLearn enables rapid exploration of the suitability of various ML algorithms for the experimental datasets. It fosters open science via transparent assessment of state-of-the-art algorithms in a standardized format for proteomics and other omics sciences.

**Graphical Abstract:** 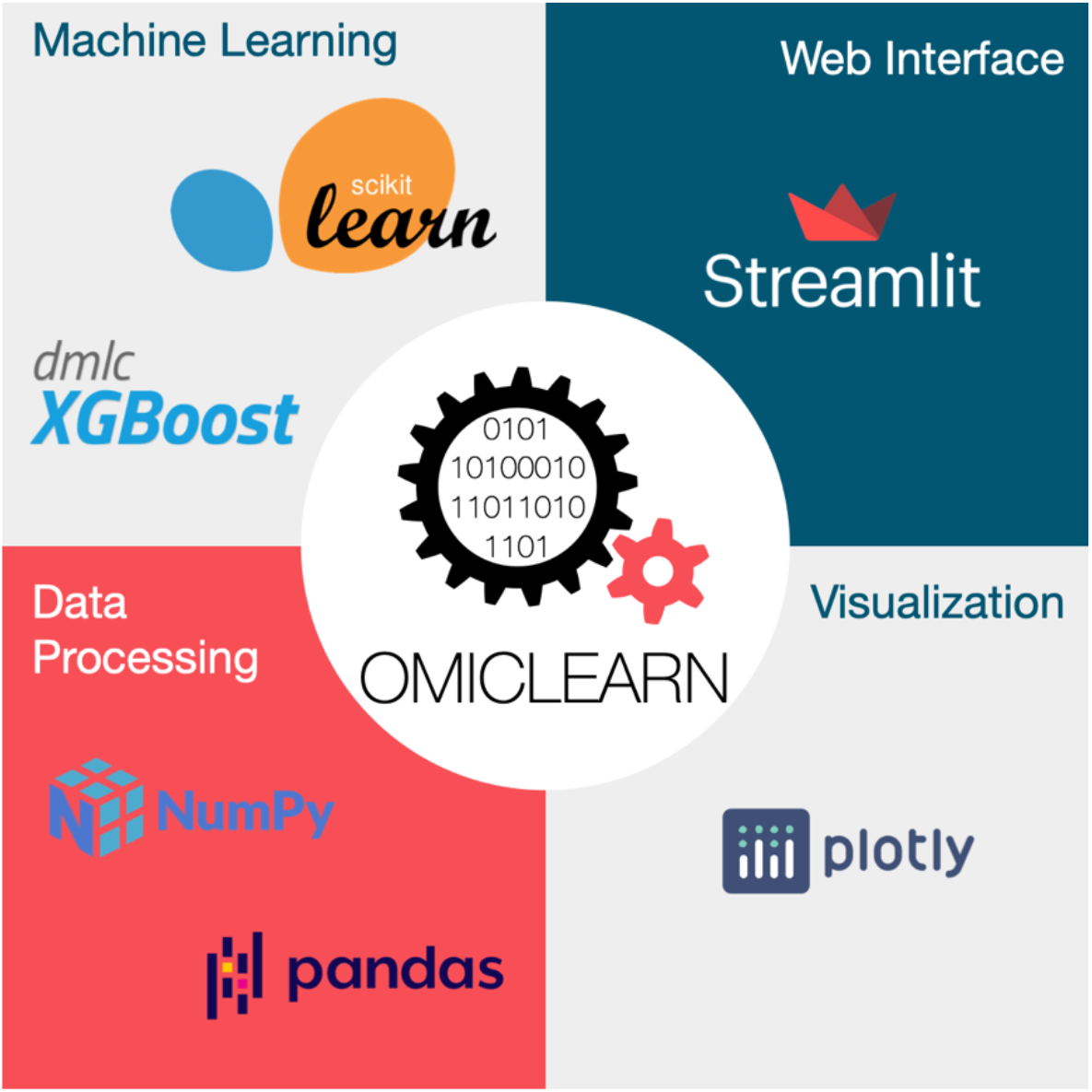

**Highlights:**

- OmicLearn is an open-source platform allows researchers to apply machine learning (ML) for biomarker discovery
- The ready-to-use structure of OmicLearn enables accessing state-of-the-art ML algorithms without requiring any prior bioinformatics knowledge
- OmicLearn’s web-based interface provides an easy-to-follow platform for classification and gaining insights into the dataset
- Several algorithms and methods for preprocessing, feature selection, classification and cross-validation of omics datasets are integrated
- All results, settings and method text can be exported in publication-ready formats

## Introduction

Machine learning (ML) is one of the most exciting opportunities for transforming scientific discovery today. While ML and its first algorithms were conceptualized decades ago, increasing computational power and larger datasets have now clearly demonstrated the superiority of ML approaches over classical statistical approaches in many applications. Concurrently, advances in omics technologies have enabled the generation of large and complex biological datasets from the analysis of hundreds to thousands of samples (Cominetti et al., 2016; Demichev et al., 2020a; Geyer et al., 2016a, 2021; Niu et al., 2020), which now allows ML to extract meaningful biological information from the data. This also applies to mass spectrometry (MS)-based proteomics, which has become the method of choice for the quantitative investigation of the entirety of proteins and their modifications in a biological system (Bache et al., 2018; Geyer et al., 2016b, 2017; Meier et al., 2018). Continuous technological advances are transforming MS-based proteomics from a basic research tool to a powerful clinical technology. As technological challenges in robustness, throughput, and reproducibility are being solved, MS-based proteomics is becoming increasingly popular for the analysis of clinical samples and an ideal tool for biomarker discovery. The development of automated sample preparation pipelines and increasingly robust liquid chromatography (LC) and MS systems enable the analysis of large studies encompassing hundreds or thousands of samples (Bache et al., 2018; Geyer et al., 2016b, 2017; Meier et al., 2018).

Large datasets are challenging to analyze in conventional ways but are well suited to ML algorithms, which can identify promising protein signatures and predict physiological states based on proteome data and additional clinical metadata. Recently, we applied ML in studies comprising hundreds of cerebrospinal fluid (CSF) or urine samples to predict the manifestation of neurodegenerative diseases (Bader et al., 2020; Virreira Winter et al., 2021). In these projects, established biomarkers associated with the investigated diseases ranked among the top candidates such as tau, SOD1 and PARK7 in Alzheimer’s Disease (AD), and VGF and ENPEP in Parkinson’s Disease (PD), and potential novel ones were uncovered.

For experimental researchers, the widespread application of ML to proteomics and other omics datasets, however, is often hampered by lacking access to the existing, best-suited ML technology. Popular packages such as scikit-learn or XGBoost allow predictive data analysis in principle, but researchers still require programming knowledge to write their own ML pipelines (Chen and Guestrin, 2016; Pedregosa et al., 2011). Additionally, the currently available packages in Python or R are not always easy-to-follow by non-specialists since they typically have no graphical interface. A noteworthy exemption is the Galaxy project, a server-based scientific workflow system that aims to make computational biology more accessible (Afgan et al., 2018). However, with more than five-hundred workflows readily available, it can be cumbersome to navigate without having domain knowledge.

Hence, access to powerful ML is restricted to experienced data scientists. Furthermore, to reproduce published results, the same software environment needs to be set up and configured with the matching package versions and random seeds. Especially in ML, selecting the appropriate methods is far from obvious to the non-specialist. Moreover, many parameters can be altered to tune the algorithms, which might change from version to version, resulting in reproducibility issues. While several packages exist that perform automatic optimization of parameters, manual verification and benchmarking of algorithms is limited. Additionally, omics sciences and ML require special domain knowledge as metrics can be deceiving and algorithms might need special preselection or preprocessing steps. For instance, in some studies, the receiver operating characteristics (ROC) curve might be useful to confirm the performance, while precision-recall (PR) curves are mandatory in imbalanced datasets (Davis and Goadrich, 2006). Thus, transparent and open-source software would be favorable, particularly in the interest of open and reproducible science (McDermott et al., 2019).

To address all these issues and to help current initiatives on biomarker discovery, we here introduce OmicLearn, a ready-to-use ML web application specifically developed for omics datasets. We describe OmicLearn’s architecture and show its benefits by applying it to a recently published proteomics study, investigating alterations in the CSF of AD patients (Bader et al., 2020). OmicLearn incorporates community efforts by building on scientific Python libraries and is available as open-source. It can be accessed via the hosted webserver or downloaded for local deployment.

## Results

### Overview of the OmicLearn architecture

Technological progress in MS-based proteomics now enables the large-scale measurement of human liquid biopsy samples such as plasma, urine or CSF. For optimal analysis of such datasets, statistical analysis methods that are capable of learning from extensive datasets such as ML algorithms are needed. ML is especially promising as it allows us to identify previously unseen dependencies and can be used to predict outcomes. To supply a ready-to-use ML interface that can be applied by experimental researchers without prior knowledge in ML, we developed OmicLearn. It is a Python-based, open-source and interactive browser application that allows the usage and parameter-tuning of multiple ML algorithms. The results are visualized by exportable graphics and charts, allowing the direct performance evaluation of the predictions. The goal of OmicLearn is to explore and benchmark how a variety of ML classifiers would perform on the data. It is not intended to export ML models to be deployed in decision making, which requires many additional scientific, statistical and regulatory considerations. That said, OmicLearn can be applied to any type of omics dataset in a tabular format and allows the export of parameters and results in a publication-ready format.

OmicLearn consists of a central web interface, an analysis core and visualization (Fig 1). Within the analysis core, data processing builds on open-source data manipulation tools such as Pandas (McKinney, 2010) and NumPy (Harris et al., 2020), which are specifically designed for multi-dimensional matrices and arrays. To implement state-of-the-art ML and preprocessing methods, we built OmicLearn on scikit-learn and advanced machine learning algorithms such as XGBoost (eXtreme Gradient Boosting). Scikit-learn is a widely used library for classification, regression, and clustering problems, which incorporates common preprocessing, feature selection and cross-validation techniques needed in ML (Chen and Guestrin, 2016; Pedregosa et al., 2011). XGBoost comes with additional algorithms, improved performance, and optimized memory usage.

**Figure 1.**
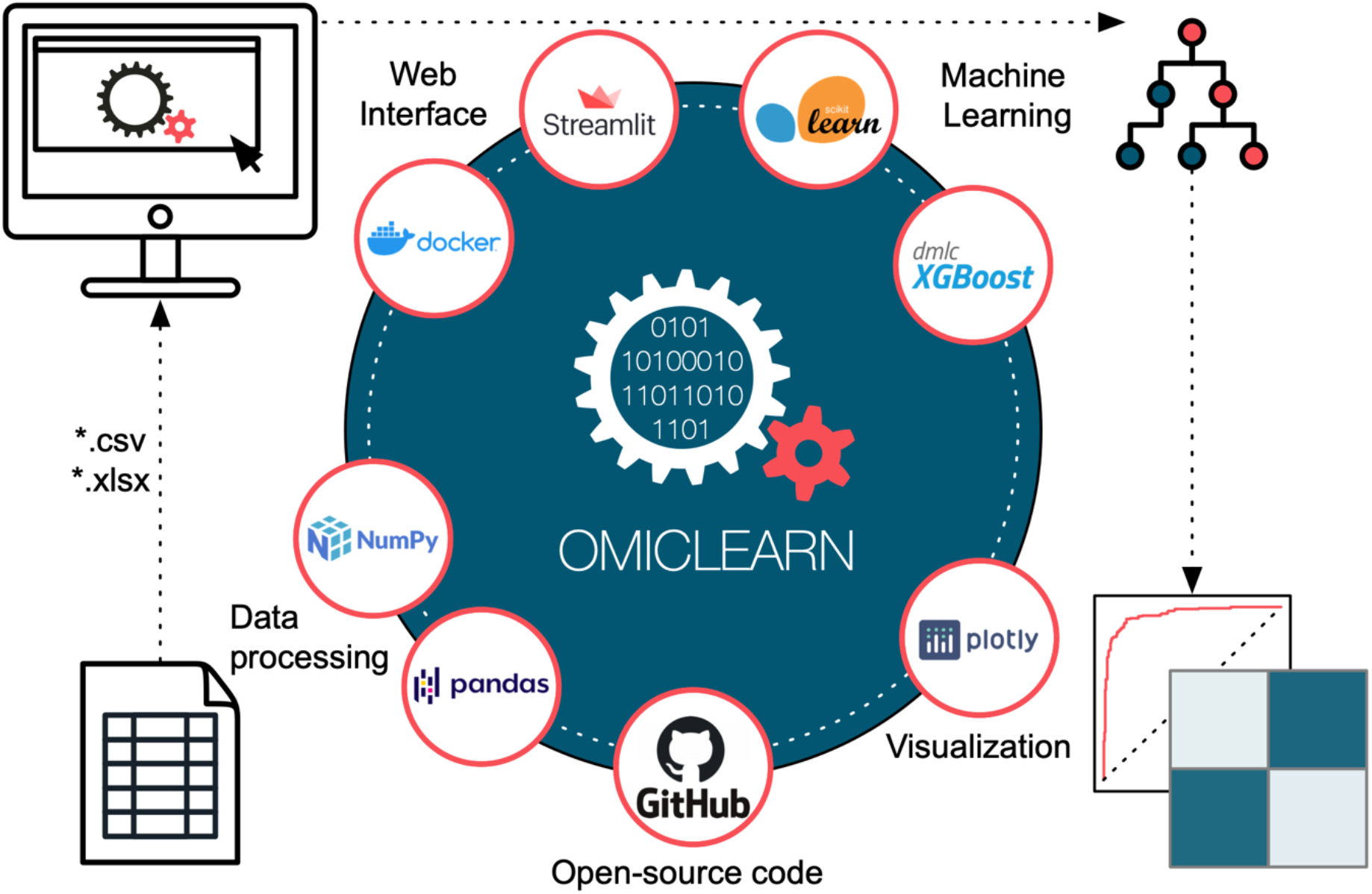
The OmicLearn architecture. *Left side*: OmicLearn is integrated in an interactive web-based tool that is built on the Streamlit package. The application is containerized via Docker, so that it can be easily deployed in a scalable cloud environment. Tabular experimental data files can be uploaded to OmicLearn as *.csv or *.xlsx (Excel format). Internally, OmicLearn uses the NumPy and Pandas packages to import and handle data. *Right side*: OmicLearn has access to the large Machine Learning libraries of scikit-learn as well as additional algorithms such as XGBoost. The pipeline is set up to perform classification tasks on omics datasets with multiple cross-validations of results. Various performance metrics are displayed, leveraging the Plotly library. The OmicLearn repository is hosted on GitHub and is open-source.

The interactive web-interface and visualization components are built on the recently developed open-source framework Streamlit (https://www.streamlit.io). Dropdown menus allow the straightforward definition of dataset-specific variables and the selection of various parameters for different ML algorithms. A core feature that facilitates usage especially for novice user is the automated interface update based on previously made selections, preventing invalid choices. Results are visualized with the graphic Python library Plotly (https://plotly.com/python) to generate high-quality interactive graphs, which can be exported as *.pdf, *.png or *.svg. For guidance, we implemented a Wiki documentation (https://github.com/OmicEra/OmicLearn/wiki) that provides background knowledge about OmicLearn, its ML algorithms, and the available methods. Additionally, the Wiki supplies information on clinical MS-based proteomics, an installation guide and a user manual for the tool.

The OmicLearn code is released as open-source under the Apache License (2.0). The tool itself is available on GitHub https://github.com/OmicEra/OmicLearn, which includes the documentation, the complete source code and the example dataset described below. OmicLearn can be installed and run locally to enable use in restricted or sensitive environments. To facilitate installation or usage in a cloud environment, we included a Dockerfile for containerization. Alternatively, a running instance of the online app can be accessed via the website https://OmicLearn.com.

### How to use OmicLearn?

Datasets can be uploaded via drag and drop or browsing a local drive. Internally, OmicLearn is built on the widely used Pandas and NumPy packages to import and store data. Datasets should be supplied in a .csv or .xlsx format, which are the typical output tables of packages such as MaxQuant or DIA-NN (Cox and Mann, 2008; Demichev et al., 2020b). The datasets need to meet distinct criteria with regards to the structure of the data matrix. Each row should correspond to a sample and each column to a feature to be used for classification and every column must have a header. Features can be supplied as two types: main and additional. Main features typically comprise the abundance information of every analyte (e.g. protein or metabolite intensities), while additional features are associated with clinical information such as age, sex or disease status of the samples or subjects. These additional features have to be supplied starting with an underscore ’_’ (e.g. ’_age’) for OmicLearn to distinguish them from main features. For quickly testing out the features of OmicLearn without uploading a custom file, we provide a tutorial sample file as well as a sample data set from a recently published study on biomarker discovery in AD using cerebrospinal fluid (CSF) (Bader et al., 2020).

Once a file is uploaded, OmicLearn’s ML interface appears, consisting of two separate selection menus for ML options and for dataset-specific feature definitions (Fig 2A). The core steps of the pipeline can be found in the left sidebar, where the user can specify individual parameters for random state, preprocessing, feature selection, classification and cross-validation. As an example, OmicLearn offers the choice between several algorithms for classification, including AdaBoost, Logistic Regression, Random Forest, XGBoost, Decision Tree, KNN Classification and linear Support Vector Classification (briefly described with additional links in the OmicLearn Wiki). Within the interactive interface, several hyperparameters can be defined according to the chosen model or algorithm. Furthermore, a random state slider allows the specification of a seed state to make random operations such as train-test splits deterministic to ensure reproducibility of the predictions.

**Figure 2.**
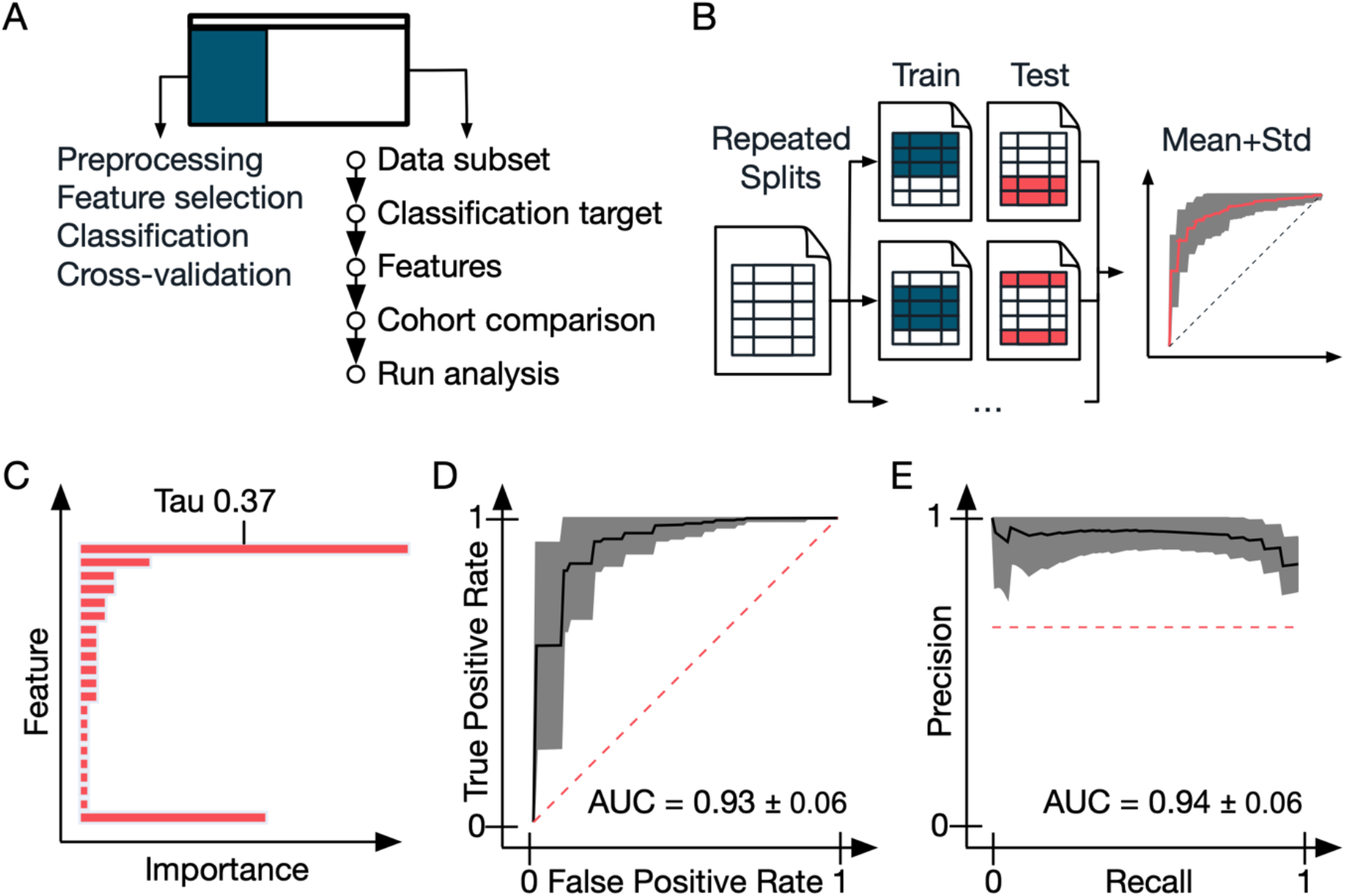
Functional flow of OmicLearn and example performance metrics. A. The OmicLearn’s landing page is comprised out of two functional elements. The left side allows setting the parameters for the ML pipeline such as selecting the ML classifier and setting algorithmic parameters. The right side allows manipulating the dataset and exploration of results. The process is interactive and follows a linear flow, e.g. whenever an option is selected only choices that will match the previously selected parameters will be shown. B. Cross-validation (CV) strategy: Data is repeatedly split into train and validation sets, so that means and standard deviations can be estimated. C. Feature importance: This plot shows the Feature importance (weights) from the ML classifier, averaged over all classification runs. The annotation on the y-axis is interactive and will directly link to NCBI. D. Interactive Receiver Operating Characteristics (ROC) curve: The ROC curve shows individual CV splits as well as an average ROC curve with a confidence interval. E. Interactive Precision-Recall (PR) curve: The PR curve shows individual CV splits as well as an average PR curve with a confidence interval.

The underlined headlines of the ML options such as ‘Feature selection’ are linked to the Wiki. Here, we supply a stepwise manual to apply OmicLearn and more information for all sections and methods. Moreover, the user will find references for supporting information for the ML algorithms, metrics and scores.

Subsets of uploaded datasets can be created based on an additional feature column, e.g., when having a multicenter study and only wanting to train on the data of a specific study center.

Within the ‘Classification target’ section, the user can specify the column that contains the classification target. Here, they can define two classes that are based on unique values within this column that the classifier will be trained to distinguish. In a typical setup, this could be the disease state to classify patient and control samples. If there are more than two unique values, each class can be defined to consist of multiple values, or values can be excluded when training the classification algorithm.

OmicLearn also allows to include additional feature columns in the classification, which can be selected under the ‘Additional features’ section. If the column contains non-numerical data such as ‘condition_a’, ‘condition_b’ and ‘condition_c’ for a category, OmicLearn will convert the values to numerical data such as 0, 1 and 2. In this section, users might upload their *.csv file (comma “,” separated), where each row corresponds to a feature to be excluded. Furthermore, it is possible to manually select the main features via ‘Manually select features’. Lastly, the option ‘Cohort comparison’ allows using one of the additional feature columns to split the dataset into different cohorts to train on one cohort and test on the other. Once all parameters are set, clicking on the ‘Run analysis’ button will initiate the selection of the best features and calculation of the predictive model.

### Interpretation of results

OmicLearn reports various metrics, ranging from reports on important features to the evaluation of the applied ML models. These results are displayed in several tables and graphs. A bar plot ranks the features with the highest contribution to the prediction model (20 in our tutorial dataset; Fig 2C) from all of the cross-validation (CV) runs. For instance, in our sample dataset analysis, the known biomarker tau (P10636) displayed the highest feature importance value, as described in the original study. This information is also available as tables in *.csv format. To comfortably retrieve more knowledge about these features, we directly linked their IDs or names to a National Center for Biotechnology Information (NCBI) search.

In order to evaluate the performance of an ML model, a study needs to be split into train, validation, and holdout (test) set. Optimization is performed using the training and validation set, and the model that is ultimately used is being evaluated using the unseen holdout set. As already mentioned, OmicLearn is intended to be an exploratory tool to assess the performance of algorithms when applied to specific datasets at hand, rather than a classification model for production. Therefore, no holdout set is used and the performance metrics have to be interpreted accordingly. This also prevents repeated analysis of the same dataset and choosing the same holdout set from leading to a selection bias and consequent over-interpretation of the model.

The strategy of splitting data is crucial to overcome the common ML problems of over- or underfitting. Overfitting occurs when applying a model with high complexity that learns on unrelated noise. Over-fitted models will be capable of describing the sample with high accuracy but will not generalize well when validating on another dataset. In our context, this is frequently observed when study-specific biases are present that are not found in future observations. Underfitting happens when the model is not sufficiently complex, and is therefore not capable of learning the subtleties of the sample characteristics, resulting in sub-optimal performance. Even though the throughput of omics sciences is rapidly increasing, the number of analyzed samples is generally small compared to the number of features that can be measured. To illustrate, a sample cohort may be in the range of hundreds but we are measuring thousands of proteins, making ML particularly prone to overfitting. In order to use the existing data most efficiently, we use CV, in which data is repeatedly split into train and validation sets (*RepeatedStratifiedKFold* method). For this purpose, we integrated a stratified splitting technique, meaning that the original class ratio will be preserved for the splits. OmicLearn offers additional split methods such as *StratifiedKFold* and *StratifiedShuffleSplit*, which can be selected in the ML options (Fig 2B). These measures prevent models scoring well, that have ‘learned’ to simply predict the majority class (i.e. always predicting that a rare condition is absent).

The number of features that are being used for the model can be either selected by the user or automatically selected with feature selection algorithms built into OmicLearn. The feature importance scores obtained from the classifier after all CV runs are displayed in a horizontal interactive bar chart and an exportable table (Fig 2C). The feature importance is additionally provided in tabular format and contains the standard deviations. The feature selection is applied for each split during the CV process so that no information leakage occurs.

We further implemented Receiver Operating Characteristics (ROC) for a graphical representation of model performance (Fig 2D). They display the true positive rate (sensitivity) against the false positive rate (1 - specificity) in an easily interpretable form. In the supplied plot, the mean ROC curve (black) is displayed together with the standard deviation (grey background) of the different curves from the various train and validation set splits. The Area Under the Curve - Receiver Operating Characteristics (AUC-ROC) is a numerical value to assess the prediction; it would be 1.0 in the case of a perfect discrimination. In addition, we use Precision-Recall (PR) curves displaying the sensitivity (recall) against the positive predictive value (Fig 2E). PR curves are valuable for performance assessments, especially in when dealing with imbalanced datasets, where one class is more frequent than the other (Saito and Rehmsmeier, 2015). To further evaluate the quality of the predictions, we supply a 2×2 ‘confusion matrix’ to compare predicted and actual classes. In the sample dataset, it displays the number of correctly predicted positive and negative AD patients as well as the number of false-positive and false-negative predictions.

The overview of all results is available in one comprehensive table in the ‘Results’ section. We further provide a publication-ready summary text for describing packages, libraries, methods, and parameters. Finally, since researchers might perform multiple runs in OmicLearn to explore different learning conditions, previous results are listed in the ‘Session History’ section. In this way, users can easily compare current with previous results. Additionally, a download option for the session history as *.csv exists. The graphics generated by OmicLearn can be saved in a publication-ready format such as *.pdf, *.svg, and *.png, and all tables are available as .csv files.

### Application examples

The underlying type of a ML classifier can have a drastic effect on the model performance depending for a given dataset it is applied to. Therefore, models should be selected to fit the nature of the problem. In the analysis of our sample AD dataset with OmicLearn, we quickly evaluated seven ML algorithms. Each model showed different performance on predicting Alzheimer’s disease status. To illustrate such effects, we use several OmicLearn’s metrics in a ‘Run results for classifier’ table and graphs to show the influence of classifiers (Fig 3A). The AUC-ROCs ranged from 0.63 to 0.93. This result cannot be due to differences other than the model as we had defined the same data subsets and other selections such as additional features and chose the same options for preprocessing, missing value imputation, feature selection and cross-validation (Fig 2A). Further investigating the individual model performance highlights interesting characteristics. While the majority of the models achieve an AUC-ROC of larger than 0.8, there are some outliers with much lower performance such as KNeighbors with 0.63 ± 0.12 and the decision tree model with 0.73 ± 0.09. Interestingly, a rather simple model (LogisticRegression) obtained an average AUC-ROC of 0.84 ± 0.09, which is higher than the more sophisticated support vector model (LinearSVC) with a mean score of 0.81 ± 0.09.

**Figure 3.**
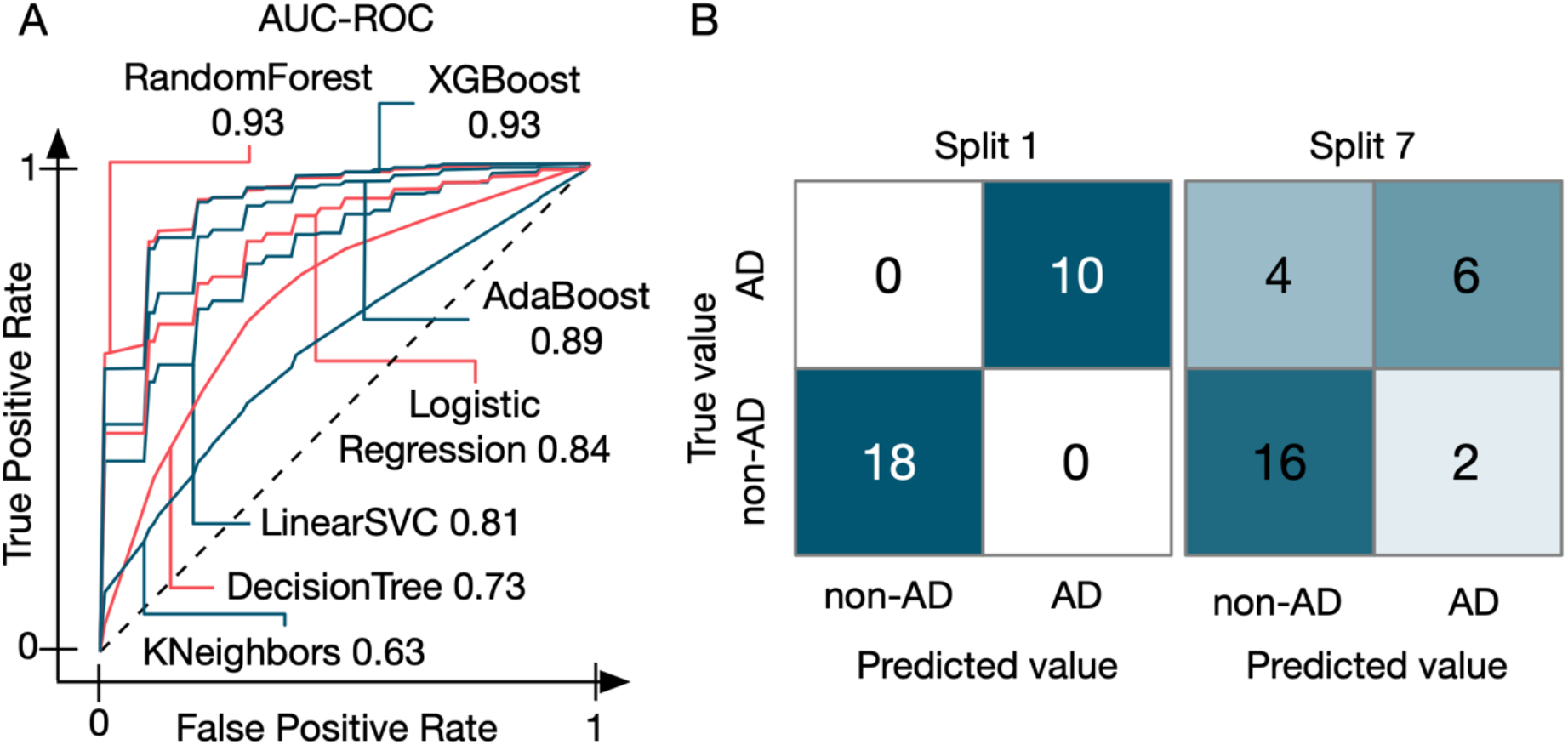
Application examples of OmicLearn. A. ROC curves generated from the AD dataset for multiple ML models. The achieved AUC-ROC ranged from 0.63 to 0.93. The different ML algorithms are indicated with their AUC-ROC value. B. Examples of two different splits of the AD dataset for one ML model. Split 1 resulted in perfect accuracy and exhibited no false classification. Split 7 of the same cross-validation run had several false classifications and hence lower performance, highlighting the importance of cross-validation.

One of the best models for this application is the XGBoost classifier, which achieves an AUC-ROC of 0.93 ± 0.06. Note that the minimum AUC-ROC for a single CV split was 0.77, while the maximum was 1. This emphasizes that repeated validation is necessary to avoid misinterpreting performance on favorable or unfavorable splits.

A confusion matrix facilitates understanding performance metrics by showing actual numbers for each class (Fig 3B). To display the individual cross-validation splits, OmicLearn provides an interactive confusion matrix with a slider for picking a split. We found even perfect splits (e.g. split 1) that classified all Alzheimer’s patients (10/10) and non-Alzheimer’s patients (18/18) correctly. In contrast to that there are splits that are much worse (e.g., split 7), which only classifies (6/10) and (16/18) correctly, highlighting the variance in prediction accuracy.

## Discussion

Recent technological advances are dramatically improving robustness, throughput and reproducibility of omics technologies such as genomics, proteomics and metabolomics. This has sparked an increasing interest in using these technologies for biomarker discovery with large cohorts of clinical samples. More generally, the analysis and interpretation of large biological datasets obtained from omics technologies are complex and require automated computational workflows. In addition to the statistical tests that are typically applied, ML is an increasingly powerful tool to extract meaningful information and to obtain a deeper understanding of the underlying biology. The application of ML algorithms to large omics datasets, however, remains a challenge in many ways. Individual ML pipelines need to be established, specialized knowledge of data scientists or bioinformaticians is required and the applied workflows often lack transparency and reproducibility. While the number of studies applying ML to omics datasets is rapidly increasing, issues associated with transparency of analyses, validation of existing results and reproducibility are increasingly recognized and a matter of concern in the field.

To make ML algorithms easily accessible and understandable for experimental researchers, we developed OmicLearn, a browser-based app that allows applying modern ML algorithms to any omics dataset uploaded in a tabular format. Although developed with clinical proteomics in mind, it is in no way limited to this application. OmicLearn offers several ways to explore the effect of a variety of parameter settings on ML performance and comes with a detailed Wiki containing background information and a user manual. Within OmicLearn, multiple methods are available for preprocessing, feature selection algorithms, classification and cross-validation steps together with hyperparameter tuning options. Furthermore, OmicLearn enables researchers to export all settings and results as publication-ready figures with an accompanying methods summary. This enables researchers to apply the identical pipeline to multiple omics datasets or reproduce existing results and simplifies the application and usage of ML algorithms to any tabular data without requiring any prior ML knowledge. With its user-friendly interface, OmicLearn enables researchers to upload a dataset with features, such as protein levels and any associated clinical information such as disease status, to train and test a model and provide new valuable insights into the dataset. OmicLearn aggregates the methods and algorithms from Python ML library scikit-learn together with XGBoost. Furthermore, it combines several best practices for preprocessing and feature selection steps to apply them to the files uploaded by users, such as MS-based proteomics datasets. To demonstrate its usability, we have applied various ML algorithms to a recently published study that investigated changes in the CSF proteome of AD patients. While we showcased our app on proteomics data, it can be applied to tabular data obtained using other omics technologies such as genomics or metabolomics. A principal challenge that remains for all ML approaches is explainability. In a biomarker discovery context, features that give highly accurate models could originate from inherent study biases so that scrutinizing results with respect to the underlying biology is imperative. The interactive nature of OmicLearn should aid in this process.

A key finding is that ML requires repeated cross-validation of results as biased splitting of data can result in drastic performance variation, which can be larger than the performance difference of different classifiers. While some models will have better performance, the baseline classification accuracy of all classifiers should be in the same range and the user should be able to achieve competitive results with OmicLearn. This also suggests that it is beneficial to stringently benchmark a study with a relatively standard model and having a good understanding of the baseline performance instead of purposely building a model for a particular study. In this way, OmicLearn also helps to democratize ML in the field as results will be more comparable and differences in model performance easier to understand.

In summary, OmicLearn is an easy-to-use powerful tool for exploratory data analysis. It gives a rapid overview of how well the supplied data perform in a classification task and can be applied to fine-tune and optimize ML models. However, on its own, it does not provide biomarker panels or models ready to be used in diagnostics. Potential improvements of OmicLearn include the diversification of ML classification algorithms and the inclusion of other sophisticated optimization and preprocessing methods such as standardization, imputation of missing values and data encoding. We have found OmicLearn to be an indispensable tool in analyzing clinical proteomics datasets and hope that it will provide similar benefits for a large community of researchers in the field of biomarker discovery.

## Author contributions

FMT and MTS wrote the code of OmicLearn, drafted its GitHub repository, and designed the figures. FMR and AV tested the application and revised the code. PEG, FMT and MTS wrote the manuscript and MTS conceived the original idea of OmicLearn. All authors performed research, analysis, manuscript writing and provided critical feedback.

## Acknowledgments

We thank Halil I. Bilgin for helpful discussions on OmicLearn. We thank our former colleagues at the Max-Planck Institute of Biochemistry, Jakob M Bader and Ozge Karayel for initial discussions related to Machine Learning.

